# Evolution of multicellularity by collective integration of spatial information

**DOI:** 10.1101/2020.02.20.957647

**Authors:** Enrico Sandro Colizzi, Renske M.A. Vroomans, Roeland M.H. Merks

## Abstract

At the origin of multicellularity, cells may have evolved aggregation in response to predation, for functional specialisation or to allow large-scale integration of environmental cues. These group-level properties emerged from the interactions between cells in a group, and determined the selection pressures experienced by these cells.

We investigate the evolution of multicellularity with an evolutionary model where cells search for resources by chemotaxis in a shallow, noisy gradient. Cells can evolve their adhesion to others in a periodically changing environment, where a cell’s fitness solely depends on its distance from the gradient source.

We show that multicellular aggregates evolve because they perform chemo-taxis more efficiently than single cells. Only when the environment changes too frequently, a unicellular state evolves which relies on cell dispersal. Both strategies prevent the invasion of the other through interference competition, creating evolutionary bi-stability. Therefore, collective behaviour can be an emergent selective driver for undifferentiated multicellularity.

## 1 Introduction

The evolution of multicellularity is a major transition in individuality, from autonomously replicating cells to groups of interdependent cells forming a higher-level of organisation [1, 2]. It has evolved independently several times across the tree of life [3, 4]. Comparative genomics suggests [5], and experimental evolution confirms [6, 7] that the increase of cell-cell adhesion drives the early evolution of (undifferentiated) multicellularity. Increased cell adhesion may be temporally limited and/or may be triggered by environmental changes (e.g. in Dictyostelids and Myxobacteria [8, 9]). Moreover, multicellular organisation may come about either by aggregation of genetically distinct cells or by incomplete separation after cell division [8, 10].

The genetic toolkit and the cellular components that allow for multicellularity - including adhesion proteins - pre-date multicellular species and are found in their unicellular relatives [8, 11–13]. Aggregates of cells can organise themselves by exploiting these old components in the new multicellular context, allowing them to perform novel functions (or to perform old functions in novel ways) that may confer some competitive advantage over single cells. Greater complexity can later evolve by coordinating the division of tasks between different cell lineages of the same organism (e.g. in the soma-germline division of labour), giving rise to embryonic development. Nevertheless, the properties of early multicellular organisms are defined by self-organised aggregate cell dynamics, and the space of possible multicellular outcomes and emergent functions resulting from such self-organisation seems large – even with limited differential adhesion and signalling between cells. However, the evolution of emergent functions as a consequence of adhesion-mediated self-organisation has received little attention to date.

Mathematical models can define under which conditions multicellularity evolves, in terms of fitness for individual cells vs. the group, or in terms of the resulting spatial and temporal organisation. The formation of early multicellular groups has been studied in the context of the evolution of cooperation: by incorporating game theoretical interactions and transient compartimentalisation [14] or the possibility of differential assortment [15], it was found that adhering groups of cooperating individuals evolve. Alternatively, costly reproductive trade-offs in a structured environment can give rise to division of labour and the formation of a higher-level proto-organism capable of self-regeneration [16]. A plethora of multicellular life-cycles can emerge by simple considerations about the ecology of the uni-cellular ancestor and the fitness benefit that cells acquire by being in groups [17]. Once multicellular clusters are established, the spatial organisation of their composing cells can play an important role in determining group-level reproduction - possibly leading to the evolution of cell-death [18] and to specific modes of fragmentation of the aggregate [19, 20] that increase overall population growth.

In these models, multicellularity is either presupposed or its selective pressure is predetermined by social dynamics, by directly increasing fitness of cells in aggregates or by adverse environmental conditions that enforce strong trade-offs. Here we investigate the origin of this selective pressure, motivated by the idea that multicellular groups emerge as a byproduct of cell self-organisation and cell-environment interactions, and subsequently alter the evolution of their composing cells. We expect that a selective pressure to aggregate can arise from the emergent functions of the multicellular group, without requiring explicit selective advantages and disadvantages for cells in a group. We therefore present a computational model of an evolving population of cells where fitness is based solely on how adequately a cell responds to a spatially and temporally heterogeneous environment, regardless of whether they belong to an aggregate.

We draw inspiration from the life cycle of the slime mould *Dictyostelium discoideum* and in particular its slug phase (described in e.g. [21]), in that we let cells move preferentially towards the source of a noisy chemotactic gradient. Cells have a higher chance to reproduce when they are close to the source of the gradient at the end of each season. Upon reproduction, cells can evolve their adhesion to one another - and therewith undifferentiated multicellularity - when the emergent collective behaviour of cell clusters turns out to be advantageous within the structure of the changing environment.

With this model setup, we consider collective cell movement as an emergent driver of multicellularity. Collective movement is important in simpler multicellular organisms [21–23] as well as in many processes within complex multicellular organisms, such as embryogenesis, tissue repair and cancer [24, 25], and has been modelled extensively [26–32]. In our model, cells perform chemotaxis towards the source of a noisy, shallow chemokine gradient. While individual cells follow the chemotactic signal very inefficiently, groups of cells exhibit efficient chemotaxis due to the “many wrongs” principle [33]. We show that this emergent property of cell groups is sufficient to select for high levels of adhesion and multicellularity, despite the fact that fitness is only defined at the cell level.

## 2 Results

### Model setup

We consider a population of cells on a grid that search for resources to be able to replicate. We explicitly account for cell shape and interactions by implementing a 2D hybrid Cellular Potts Model on a square lattice [34–36]. Cells can adhere to each other if they express matching ligands and receptors on their surface. A better ligand-receptor match is translated to stronger adhesion, quantified by cell-cell and cell-medium adhesion energy (respectively *J*_c,c_ and *J*_c,m_ in units of energy per unit surface, see Fig. 1a and Methods).

**Figure 1:**
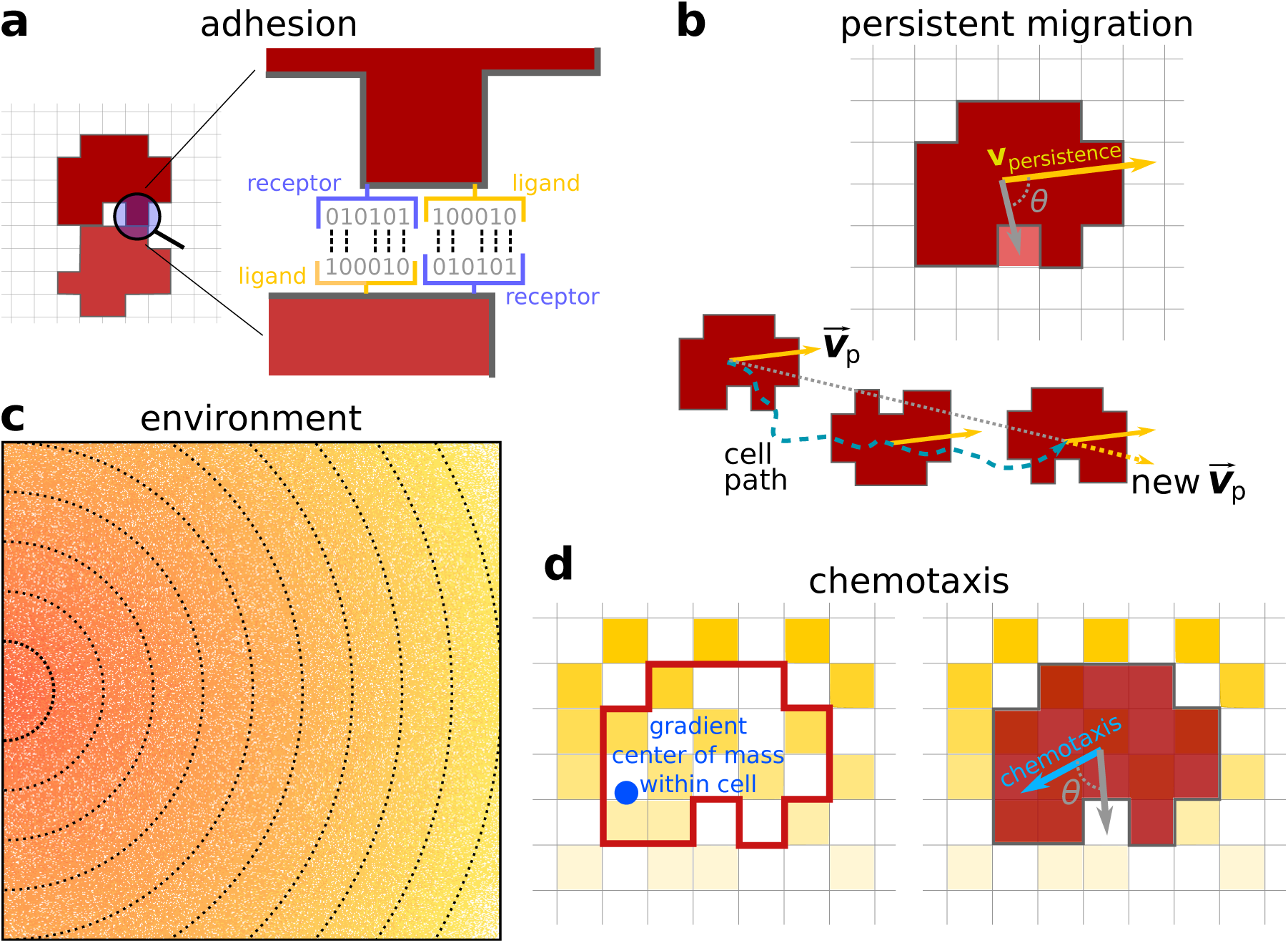
Model description. **a)** Adhesion between two cells is mediated by receptors and ligands (represented by a bitstring, see [37]). The receptor of one cell is matched to the ligand of the other cell and vice versa. The more complementary the receptors and ligands are, the lower the J values and the stronger the adhesion between the cells. **b)** Persistent migration is implemented by endowing each cell with a preferred direction of motion 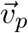. Every *τ*_*p*_ MCS, this direction is updated with a cell’s actual direction of motion in that period. **c)** The chemokine gradient in the lattice. The lines and colour indicate equal amounts of chemokine. Note the scattered empty pixels. **d)** A cell can sense the chemokine in the lattice sites that correspond to its own location. The cell will then move preferentially in the direction of perceived higher concentration, the chemotaxis vector. This vector points from the cell’s center of mass to the center of mass of the chemokine detected by the cell (the blue dot).

We implemented two types of cell migration: persistent random walk (Fig. 1b) and chemotaxis towards higher local concentrations of a chemokine (Fig. 1c). These chemokines are released by resources present at one end of the grid, creating a shallow and noisy gradient throughout the grid (Fig. 1c). Because of the noise in the gradient, the direction of cell’s chemotaxis may be different from the correct direction of the gradient. We used this model setup to assess the properties of single-cell vs. collective migration.

To explore the evolutionary dynamics of a population of cells, we let the location of the resources change seasonally (thus creating an additional temporal variation in the direction of the gradient every *τ*_*s*_ MCS). This allows cells to evolve their adhesion strength. During each season in the evolutionary simulations (i.e. one period of *τ*_*s*_ MCS) cells move due to chemotaxis and persistent migration, and may adhere to or repel each other by means of the receptors and ligands expressed on their surface (Fig. 2). At the end of the season, cells divide with a probability proportional to their distance to the peak of the gradient, thus assuming that more nutrients are present where the chemotactic signal is larger. The daughter cells inherit mutated copies of the ligand and receptor, so that their adhesive properties change with respect to the parent; cell size *A*_*T*_, strength of chemotaxis *µ*_*χ*_ and migration persistence *µ*_p_ do not evolve. Finally, we keep the population size constant by randomly culling the population in excess, at which point the new season begins. We do not select for multicellularity directly: fitness is defined at the level of the single cell, and we do not explicitly incorporate a fitness advantage or disadvantage for the multicellular state. Therefore, multicellular clusters can arise only because they perform an emergent task that single cells cannot perform.

**Figure 2:**
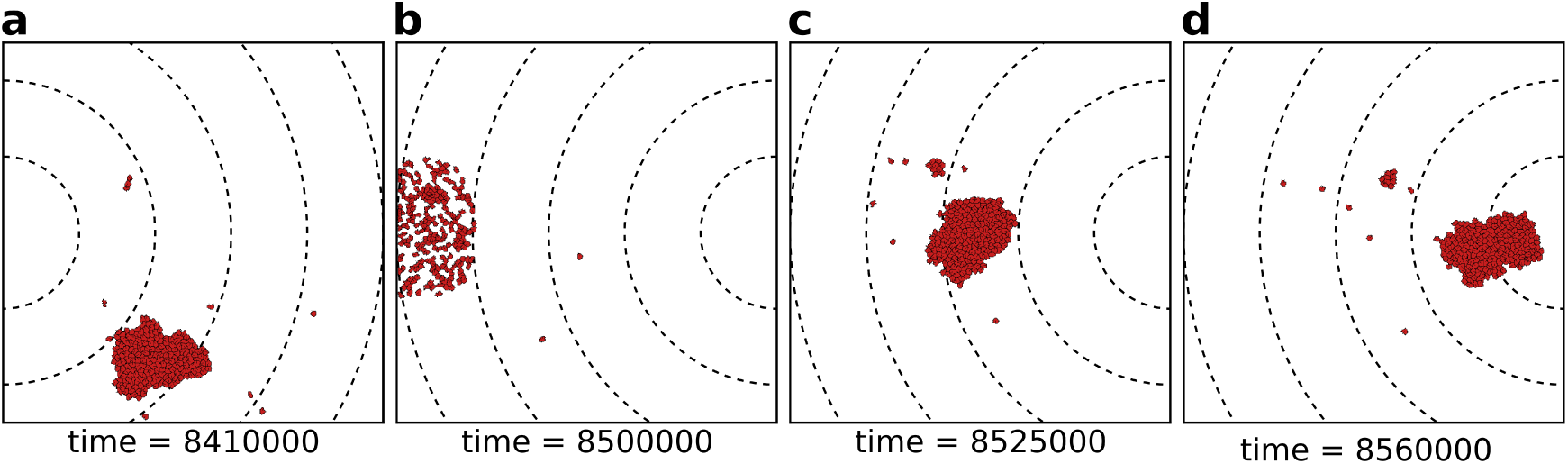
The eco-evolutionary setup of the model: **a)** A population of *N* = 200 cells moves by chemotaxis towards the peak of the gradient, which in this season is located at the left boundary of the grid. **b)** At the end of the season, cells divide, the population excess is killed randomly, and the direction of the chemotactic signal is changed, after which the new season begins (**c, d**). The snapshots are taken at the indicated time points from a simulation where a season lasts *τ*_*s*_ = 100×10^3^ MCS. Dashed lines in the snapshots are gradient isoclines.

### Strongly adhering cells perform efficient collective chemotaxis

We first assessed how well groups of cells with different adhesion strengths could reach the source of the chemotactic signal; we characterise adhesion strength by the cell’s surface tension *γ* = *J*_c,m_ − *J*_c,c_/2, so that cells adhere to one another if *γ* > 0. We recorded the travel distance of a group of cells over a fixed amount of time and compare it to the travel distance of single cells, by measuring both the position of the center of mass of the group (fig. 3a) and the position of the cell closest to the peak of the gradient (Fig. 3b). Single cells perform chemotaxis inefficiently (Fig. 3a) and show large variance between different simulations (Fig. 3b). A group of adhering cells (*γ* > 0) can migrate up the same gradient more accurately: the center of mass of this group takes much less time than single cells do to reach the peak of the gradient (Fig. 3a). Groups of cells can also perform collective chemo-taxis when they do not adhere (*γ* < 0), and when they do not have a preference for medium or cells (*γ* ≈ 0), although with lower efficiency in both cases. The velocity of the cell closest to the peak of the gradient is, instead, roughly the same regardless of adhesion strength (Fig. 3b).

**Figure 3:**
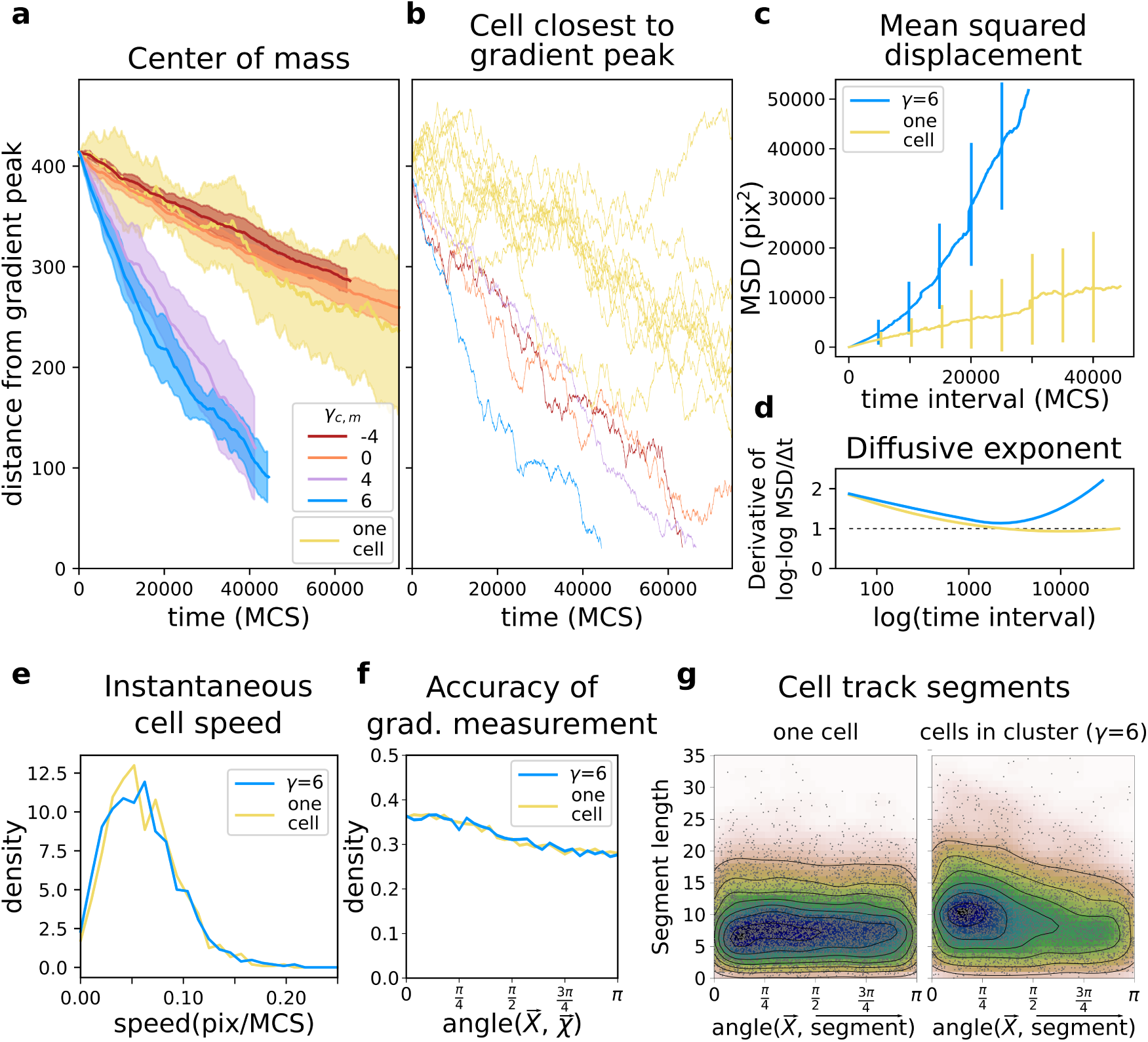
A group of cells performs chemotaxis efficiently in a noisy shallow gradient. **a)** Distance of the center of mass of *N* = 50 cells from the peak of the gradient as a function of time, for different values of *γ* ∈ [−4, 6] (five independent runs for each value), together with the average position of 10 single cells (independently run). **b)** The position of the cell closest to the gradient origin as a function of time (taken from the same simulations as in **a**), and the positions of 10 individuals cells (whose average generates the corresponding plot in **a**). **c)** Mean square displacement per time interval for two datasets each with 50 simulations of either single cells or clusters of strongly adhering cells (*N* = 50, *γ* = 6), in which case we extracted one cell per simulation. These data sets were also used for the following plots. **d)** Diffusive exponent extracted from the MSD plot, obtained from the log-log transformed MSD plots by fitting a smoothing function and taking its derivative. **e)** Distribution of instantaneous cell speeds **f)** Distribution of angles between cells’ measurement of the gradient, and the actual direction of the gradient. **g)** The length of straight segments in cell tracks vs. their angle with the actual gradient direction. Each point represents one segment of a cell’s trajectory. To extract these straight segments a simple algorithm was used (Supp. Mat. 3). Contour lines indicate density of data points.

Adhering cells have large chemotactic persistence - as shown by the super-linear shape of the mean square displacement (MSD) plot (Fig. 3c, *γ* = 6) and by a diffusive exponent consistently larger than 1 (Fig. 3d; the diffusive exponent is obtained as the derivative of the log-log transformed MSD/time curve). Instead, the MSD of individual cells (Fig. 3c, *γ* = −4) is approximately linear and their diffusive exponent tends to 1, indicating that cells’ movement is much more dominated by diffusion. Interestingly, there is no difference in the instantaneous speed of cells when they are in a cluster or when they are alone (Fig. 3e), so the higher rate of displacement of a group of adhering cells is only due to larger persistence in the direction of motion.

Within a group of adhering cells, small clusters align, pull and push on each other, generating extensions, retractions and rotations (see Supp. Video 1), so that the entire cluster visually resembles a single amoeboid cell (Supp. Video 2). This behaviour is not influenced by the presence of the chemotactic signal, since the flow field is identical in the two cases (Supp. Section S1). It is caused by the strong persistent motion of the individual cells, which aids in speeding up the movement of the cluster, but is not strictly required for chemotaxis (Supp. Section S2). Fig. 4 shows the movement of a cluster of strongly adhering cells (*γ* = 6) compared to the movement of a single cell, over the typical setup of the simulation system. Although the cluster moves straight towards the source of the gradient, individual cells follow noisy trajectories.

**Figure 4:**
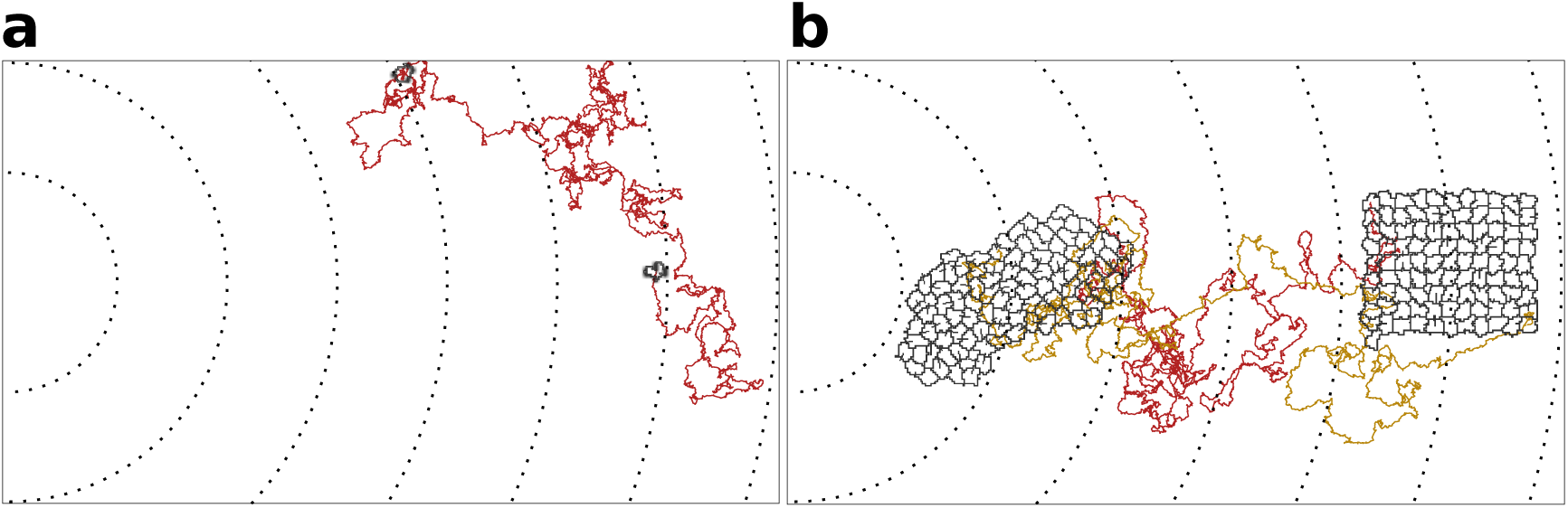
**a)** The movement of a single cell; **b)** Typical movement of a cluster of strongly adhering cells, and of the cells inside the cluster. Cells are placed on the right of the field and move towards higher concentration of the gradient (to the left of the field). Dashed lines are gradient isoclines.

We calculated the deviation of each individual cell’s measurement of the gradient as the angle 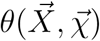 between the true direction of the gradient 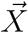 and the direction of the gradient locally measured by the cells 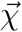 (so that 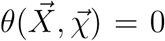 is a perfect measure). We found that the measurements of individual cells deviate significantly from the true direction of the gradient (Fig. 3e). Despite this, they are carried in the right direction by the other cells.

Altogether, collective chemotaxis of a cluster of adhering cells emerges, despite each cell measuring the gradient as poorly as cells alone (Fig. 4f). Thus, cells within a cluster must be altering each others’ paths by exerting pushing and pulling forces. To assess how these forces alter the short-timescale trajectories of cells, we extracted the straight segments from the cell tracks and assessed both the length of these segments and their orientation with respect to the gradient source (Supp. Section S3). We find that cells in a cluster tend to migrate for longer in straight lines, and that these straight lines are also more likely to be oriented towards the source of the gradient (Fig. 4g). For single cells, there is no such bias. Single cells could also improve their ability to sense the gradient by becoming bigger, since they will perceive a larger area of the chemotactic signal (Supp. Section S4). However, there are many factors that restrict how big a cell can be, such as the complexity of the metabolism and cellular mechanisms such as cell division [38, 39]. This also puts a limit on the area of a gradient that a cell can cover by sheer size. We therefore assume that cells can evolve adhesion, but not cell size.

### The evolution of uni- or multicellular strategies depends on environmental stability

The emergence of reliable chemotactic behaviour in adhering cell clusters suggests an evolutionary path to multicellularity: a population of cells may aggregate if chemotaxis is necessary for locating a resource. We therefore let cells’ adhesion - i.e. the receptor and ligands expressed by the cells - evolve in response to a seasonally changing environment. Cells closer to the peak of the gradient have a higher chance to reproduce at the end of the season (see also model setup and Methods). The receptors and ligands of the initial population are chosen such that cells neither adhere nor repel each other (*γ* = 0).

When the season lasts *τ* = 100 × 10^3^ MCS, the average adhesion between cells readily increases after only few generations (Fig. 5a): *J*_cell,cell_ decreases and *J*_cell,medium_ increases (see also Supp. Video 3 and Fig. 2 for snapshots). Fig. 5b shows that two evolutionary steady states are possible, depending on the duration of the season *τ*_*s*_. For *τ*_*s*_ < 20 × 10^3^ MCS, cells evolve to become unicellular, as cell-cell interactions are characterised by strong repulsion (*γ* < 0). Fig. 5c suggests that by selecting for *γ* < 0 cells scatter efficiently throughout the grid. Although repelling cells follow the chemotactic signal only weakly, the spreading ensures that at least some cells end up close to the source of the gradient at the end of the season. In contrast, a cluster of adhering cells would not have enough time to reach the source of the chemotactic signal when seasons are short, and would all have the same (low) fitness (see the speed of a cluster in Fig. 3). For *τ*_*s*_ > 40 × 10^3^ MCS, instead, cells evolve to adhere to one another, i.e. *γ* > 0 (see Fig. 5c for a snapshot). When seasons are sufficiently long, clusters of adhering cells have enough time to reach the source of the gradient. At this point, the fitness of cells within a cluster outweighs that of non adhering cells, because clustering increases the chances of reaching the peak of the gradient. Finally, for intermediate season duration, 20 × 10^3^ ≤ *τ*_*s*_ ≤ 40 × 10^3^ MCS, both repulsion and adhesion are evolutionary (meta) stable strategies, and the outcome of the simulation depends on initial conditions (for *τ*_*s*_ = 20 × 10^3^ MCS, the steady state with *γ* > 0 is very weakly stable).

**Figure 5:**
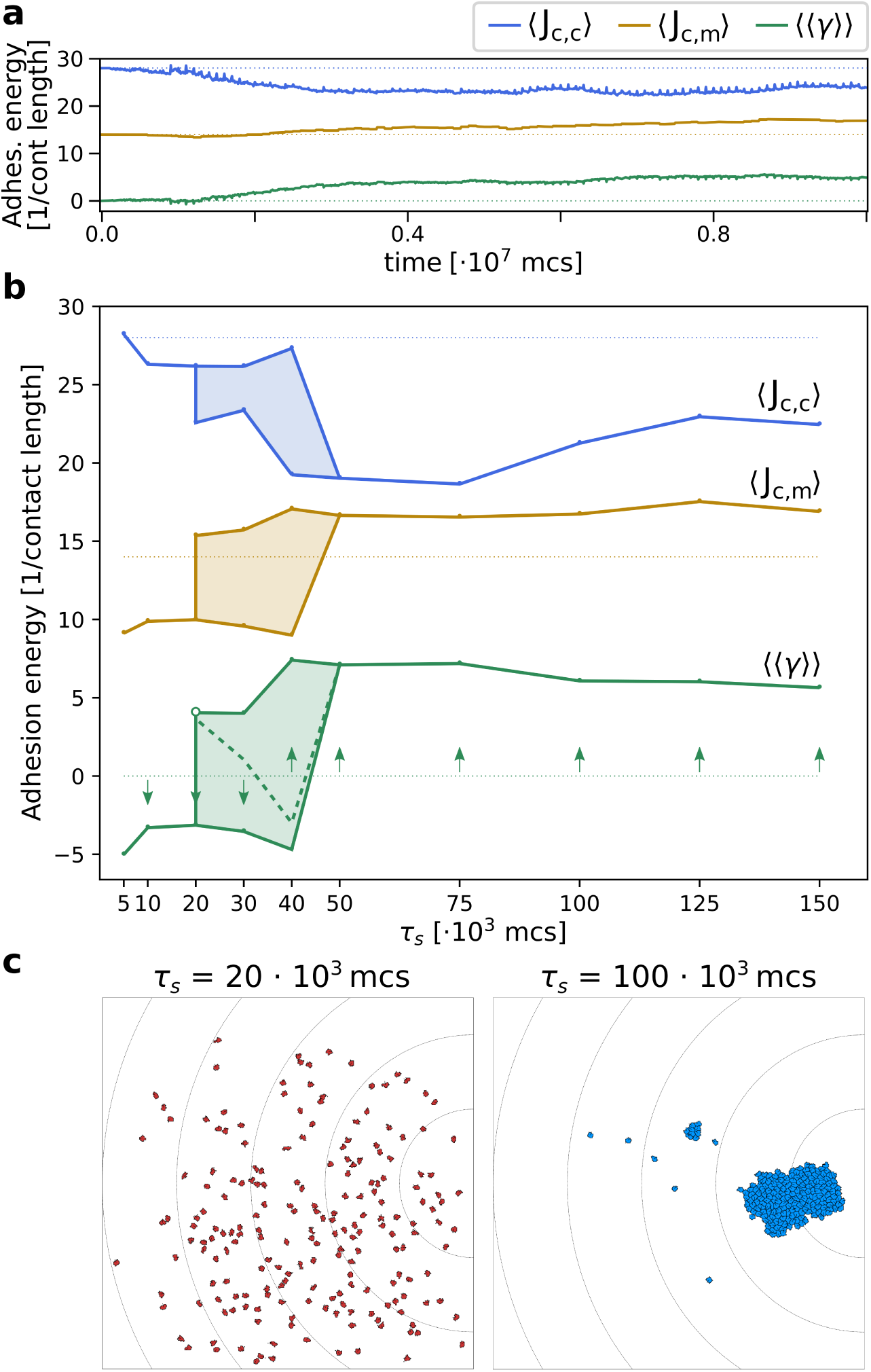
The evolution of multicellularity. **a)** Multicellularity (*γ* > 0) rapidly evolves in a population of *N* = 200 cells with *τ*_*s*_ = 10^5^ (dotted lines represent initial conditions). **b)** Multicellularity only evolves when seasons are sufficiently long *τ*_*s*_ ≥ 50 ∗ 10^3^; unicellular strategies evolve when seasons are short *τ*_*s*_ ≤ 10 ∗ 10^3^, and both strategies are viable depending on initial conditions for intermediate values of *τ*_*s*_ (the dashed line indicates the separatrix between the basins of attraction of the two evolutionary steady states). **c)** Snapshots of the spatial distribution of the population at evolutionary steady state for *τ*_*s*_ = 20 · 10^3^ and *τ*_*s*_ = 100 · 10^3^ MCS. In both panes, ⟨⟨*γ*⟩⟩ is estimated as ⟨*J*_*c,m*_⟩ − ⟨*J*_*c,c*_⟩/2, and the initial J values indicated by the dotted lines are such that *γ* = 0.

### Interference competition between unicellular and multicellular strategies causes evolutionary bi-stability

We next investigated what causes the evolutionary bistability in adhesion strategies for season duration 20 × 10^3^ ≤ *τ*_*s*_ ≤ 40 × 10^3^ MCS. We performed competition experiments between two populations of cells, one adhering (*γ* = 6) and one repelling (*γ* = −4), to determine whether a strategy can invade in a population of cells using the other strategy. We first studied how the initial distribution of the two populations affects fitness after one season of 30 × 10^3^ MCS, when both populations have equal size *N* = 100 cells. We compared a situation where the adhering cells are positioned in front of the repelling ones (Fig. 6a), with a situation where the positions of the two clusters is swapped (Fig. 6b). The distance from the peak at the end of the season (the fitness criterion) of a cluster of adhering cells is larger if they are hindered by a population of repelling cells in front of them. Next we incorporated in the competition experiments the fact that mutants invading a resident population are in small numbers and furthest away from the new peak (because they are likely born from cells that replicate most, i.e. those closest to the previous location of the peak). We simulated repelling mutants invading adhering cells by placing a large cluster of adhering cells in front of a small group of repelling ones (Fig. 6c), and conversely, a small cluster of adhering cells invading a large group of repelling cells (Fig. 6d). In both cases, the resident population physically excludes the invading one from the path to resources, and thus the distance travelled by the invading population is limited. Taken together, these results show that there is interference competition (i.e. direct competition due to displacement) between populations of cells with different strategies, which explains why the two strategies are meta-stable for intermediate season duration. This result may also provide a simple explanation for the fact that many unicellular organisms do not evolve multicellularity despite possessing the necessary adhesion proteins. Moreover, evolutionary bi-stability protects the multicellular strategy from evolutionary reversal to unicellularity over a large range of environmental conditions.

**Figure 6:**
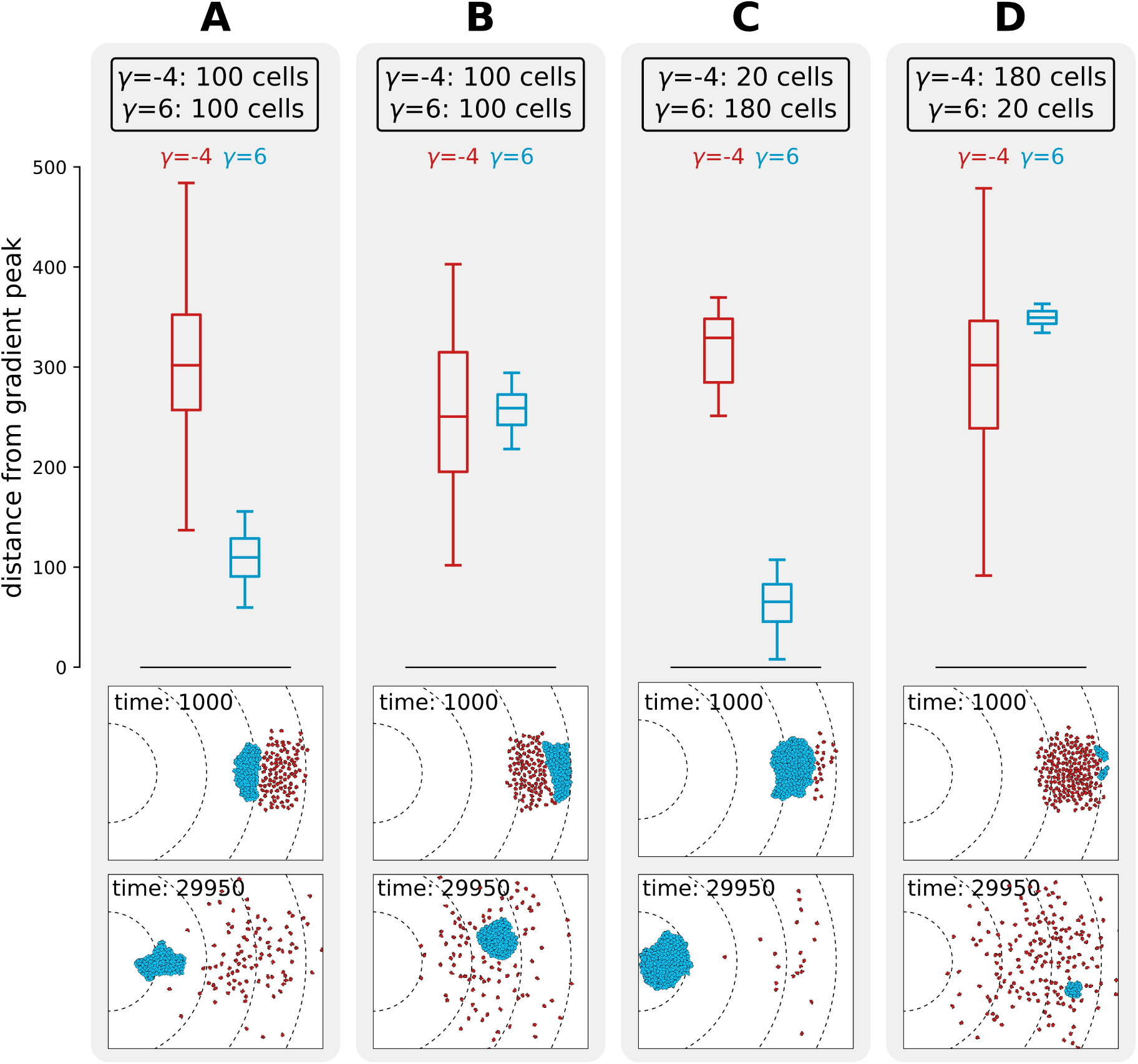
Interference competition between adhering and repelling cells explains evolutionary bistability. We let a simulation run for *τ*_*s*_ = 30 × 10^3^ MCS and then record the distance from the peak of the gradient, for two different populations of cells one repelling (in red, *γ* = −4) and one adhering (in blue, *γ* = 6), for different initial conditions. The snapshots underneath are the initial and final spatial configurations of the cells on the grid. **A)** 100 adhering and 100 repelling cells, placed so that the adhesive ones are closer to the source of the gradient; **B)** 100 adhering and 100 repelling cells, placed so that the repelling cells are closer to the source of the gradient; **C)** 180 adhering and 20 repelling cells, placed so that the adhering cells are closer to the source of the gradient; **D)** 20 adhering and 180 repelling cells, placed so that the repelling cells are closer to the source of the gradient. Dashed lines in the snapshots are gradient isoclines.

## 3 Discussion

We demonstrated that undifferentiated multicellularity can evolve in a cell-based model as a byproduct of an emergent collective integration of noisy spatial cues. Previous computational models have shown that multicellularity can be selected by reducing the death rate of cells in a cluster [17, 19], through social interaction [15, 40], by incorporating trade-offs between fitness and functional specialisation [41] or by allowing cells to exclude non-cooperating cells [42]. In these studies, direct selection for forming groups is incorporated by conferring higher fitness to the members of a cluster.

Earlier work found that multicellular structuring can emerge without direct selection when cells are destabilised by their internal molecular dynamics (e.g. the cell cycle) [43], or because of a toxic external environment [16]. In both cases, cell differentiation stabilises cell growth and arises as a consequence of physiological or metabolic trade-offs. Our work bears some similarity with these models because we do not explicitly incorporate a fitness benefit (or disadvantage) for being in a group. Our results also show that division of labour - although important is not a strict requirement for emergent aggregation. Furthermore, the simple nature of our model makes our results easily testable *in vitro*.

In many ways, the evolution of multicellularity can be compared to the evolution of collective dynamics. Previous studies on the evolution of herding behaviour showed that aggregating strategies evolve in response to highly clumped food even though the pack explores the space slowly and inefficiently before finding food [44]. In our case, aggregation leads to a highly efficient search strategy, guided by long-range, albeit noisy, gradients. Moreover, modelling cells with an explicit shape and size (something largely neglected in models of multicellularity) allows for spatial self-organisation and generates interference competition between the unicellular and multicellular search strategies. The ensuing evolutionary bi-stability stabilises unicellularity despite these cells possessing the surface protein toolkit to adhere to each other, and prevents multicellular organisation from evolutionary reversal into single cells (over a range of environmental conditions). The “automatic” outcome of spatial self-organisation provides an initial, non-genetic robustness, which can be further stabilised by later adaptations [45].

The driver for the evolution of adhesion in our model is (emergent) collective chemotaxis. This is reminiscent of the aggregate phase of the life cycle of *Dictyostelium discoideum* [21], in that a cluster of cells moves directionally as a unit following light or temperature, while individual cells are incapable of identifying the correct direction of motion. There are some important differences between our model and *Dictyostelium*, however. Individual cells are able to sense the chemotactic signal in our model, albeit inefficiently, and information about the direction of the gradient is transmitted mechanically within cell clusters. In *Dictyostelium*, individual cells cannot perceive light and thermal cues: photo- and thermo-taxis are coordinated by waves of cAMP secretion that travel through the slug. The lack of extra chemical cues to organise movement within a cell cluster in our model makes for a simpler scenario without large-scale transmission of information throughout the aggregate. Nevertheless, computational modelling has shown that long-range chemical signaling coupled to cells’ differential adhesion suffices to reproduce *Dictyostelium*’s life-cycle [26, 46]. Combining that with our evolutionary framework would likely enrich our understanding of *Dictyostelium* evolution towards partial multicellularity.

Our model of collective movement is an example of the “many wrongs” principle [33]: the direction error of each cell is corrected by the interactions with the other cells in the cluster. We adopted the Cellular Potts Model to model cells because it allowed for a straightforward implementation of the evolvable receptor-ligand system. Several other models of cell clusters and collective chemotaxis have been proposed ([30, 32]), in some cases displaying chemotaxis in qualitatively different ways (for instance without sensing the chemokine gradient, only its concentration [47]). We hypothesise that the evolutionary mechanism described here are independent of the particular cell model choice, and thus would also work with other models discussed in [30], provided that cells were able to polarise or move also in the absence of other cells.

The importance of a bottom-up approach to study the evolution of multicellularity has been repeatedly emphasised [48, 49], and a broader understanding of cells self-organisation and evolution may have applications to clinically relevant multiscale evolutionary problems, such as the evolution of collective metastatic migration of cancer cells [50–53]. Our work highlights that the properties of single cells emergently give rise to novel properties of cell clusters. These novel properties - in a downward causative direction - generate the selection pressure to form the first undifferentiated multicellular groups.

## 4 Model

We model an evolving population of cells that migrate and perform chemotaxis on a 2-dimensional lattice. Cell-cell interactions and movements are modelled with the Cellular Potts Model (CPM) [34, 35]. The evolutionary dynamics (mutations and selection) are implemented assuming constant population size. Cells undergo fitness-dependent reproduction after every season which lasts *τ*_*s*_ Monte Carlo Steps of the CPM algorithm, and then the population is culled back to its original size. After this, environmental conditions are changed and a new season begins.

### 4.1 Cell dynamics

The model is a hybrid Cellular Potts Model implemented with the Tissue Simulation Toolkit [36]. A population of *N* cells exists on a regular square lattice Λ_1_ ⊂ ℤ^2^. The chemotactic signal is located on a second plane Λ_2_, of the same size and spacing as Λ_1_. A cell *c* consists of the set of (usually connected) lattice sites 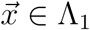 to which the same spin *s* is assigned, i.e. 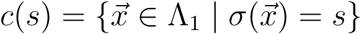.

The time dynamics are modelled as a Monte Carlo simulation. The algorithm attempts to copy the spin value 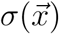 of a randomly chosen lattice site 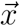 to a site 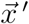 in its Moore neighbourhood. One Monte Carlo Step (MCS) consists of *L* attempted copying events, with *V* ^2^ = |Λ_1_| (the size of the lattice, and *V* one of its dimensions on a regular square lattice). Whether an attempted spin copy is accepted depends on the contribution of several terms to the energy *H* of the system, as well as other biases *Y* (explained in detail below). A copy is always accepted if energy is dissipated, i.e. if Δ*H* + *Y* < 0 (with Δ*H* = *H*_after copy_ − *H*_before copy_), and may be accepted if Δ*H* + *Y* ≥ 0 because of “thermal” fluctuations following a Boltzmann distribution:

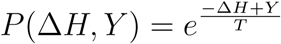

with *T* > 0 the temperature of the system, determining the probability of the fluctuations. The Hamiltonian *H* of the system consists of two terms, corresponding to adhesion and size constraint:

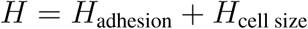

The copy biases, or “work terms”, *Y* consist of terms corresponding to cell migration and chemotaxis:

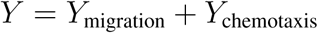

#### Cell adhesion

Adhesion between cells and to medium contribute to the Hamiltonian as:

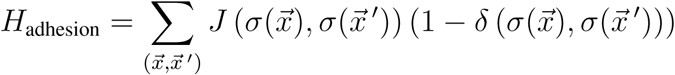

where the sum is carried out over all the neighbour pairs 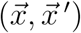, and 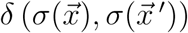 is the Kronecker delta which restricts the energy calculations to the interface between two cells, or a cell and medium.

In order to calculate the values of 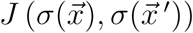, we assume that cells express ligand and receptor proteins on their surface. Ligands and receptors are modelled as binary strings of fixed length *ν* (Fig. 1, inspired by [37]). Two cells adhere more strongly (experience lower *J* values) when their receptors *R* and ligands *L* are more complementary, i.e. when the Hamming distance 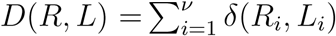 between them is larger. Thus, given two cells with spin values *σ*_1_ and *σ*_2_ and their corresponding pairs of receptors and ligands (*R*(*σ*_1_), *L*(*σ*_1_)) and (*R*(*σ*_2_), *L*(*σ*_2_)):

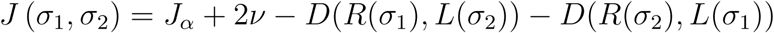

with *J*_*α*_ = 4 chosen so that the final calculation yields values for *J* (*σ*_1_, *σ*_2_) in the interval [4, 52].

Adhesion of a cell with medium is assumed to depend only on the cell (the medium is inert), and in particular it depends only on a subset of the ligand proteins of a cell. This subset consists of the substring of *L* which begins at the initial position of *L* and has length *ν*′. The value of *J* (*σ*_1_, *σ*_medium_) is calculated as:

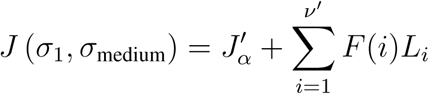

with 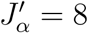 and *F*(*i*) a piece-wise defined function (a lookup table). The *J* values range in the interval [8, 20].

Encoding the energy values for cell adhesion in terms of receptor-ligand binding allows for some flexibility and redundancy. Two cells that have the same receptors and ligands (i.e. given *R*(*σ*_1_), *L*(*σ*_1_) and *R*(*σ*_2_), *L*(*σ*_2_) with *R*(*σ*_1_) = *R*(*σ*_2_) and *L*(*σ*_1_) = *L*(*σ*_2_)) can adhere with any strength (or not at all), by virtue of the particular receptor and ligand combination. Finally, implementing receptors and ligands in terms of binary strings allows for a simple evolutionary scheme, where mutations consist of random bit-flipping (more on this below). The numerical values of the various constants are chosen with two criteria in mind: the receptor-ligand system has to be long enough that many different combinations are possible, so that its evolution is more open ended; and two cells with random receptors and ligands do not (on average) adhere preferentially to each other or to the medium.

#### Cell size constraint

Cell size *A*(*c*) = |*c*(*s*)|, the number of lattice sites that compose a cell, is assumed to remain close to a target size *A*_*T*_ (equal for all cells). This is achieved by adding an energy constraint in the Hamiltonian that penalises cell sizes that are much larger or smaller than *A*_*T*_:

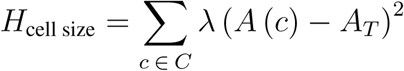

with *C* the set of cells and *λ* a scaling factor for cell stiffness.

#### Cell migration

We assume that each cell *c* ∈ *C* preferentially migrates towards a target direction given by a vector 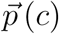 (following [54]). Lattice site copies of a cell are energetically more favourable when they are closer to the direction of that cell’s 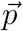:

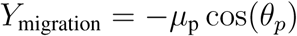

Where *µ*_p_ is the maximum energy contribution given by migration, and *θ*_*p*_ is the angle between 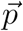 and the vector that extends from the center of mass of the cell to the lattice site into which copying is attempted. Every *τ*_*p*_ MCS the vector 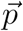 is updated: its new value is the vector corresponding to the actual direction of motion of the cell over the past *τ*_*p*_ MCS (scaled to unit) (Fig. 1). Note that all cells have the same *τ*_*p*_, but their initial moment of updating is randomised so that they do not update all at the same time.

#### Chemotaxis

Individual cells are able to migrate towards the perceived direction of a chemokine gradient. The slope of the gradient is very shallow, making it difficult to perceive the direction over the typical length of a cell. Moreover, several sources of noise are introduced: cell’s sampling error due to small size, noise due to integer approximation, and noise due to random absence of the signal.

The chemotactic signal is implemented as a collection of integer values on a second two dimensional lattice (Λ_2_ ⊂ ℤ^2^, with the same dimensions as the CPM lattice). The (non-negative) value of a lattice site represents the local amount of chemotactic gradient. This value remains constant for the duration of one season (*τ*_*s*_ MCS). The amount of chemotactic signal *χ* is largest at the peak, which is located at the center of one of the lattice boundaries, and from there decays linearly in all directions, forming a gradient: *χ*(*d*) = 1 + (*k*_*χ*_/100)(*V* − *d*), where *k*_*χ*_ is a scaling constant, *d* is the Euclidean distance of a lattice site from the peak of the gradient, and *V* is the distance between the source of the gradient and the opposite lattice boundary; 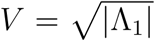 for a square lattice. Non integer values of *χ* are changed to ⌊*χ*⌋ (the smallest integer larger than *χ*) with probability equal to ⌊*χ*⌋ − *χ*, otherwise they are truncated to ⌊*χ*⌋ (the largest integer smaller than *χ*). Moreover, the value of *χ* is set to zero with probability *p*_*χ=0*_ to create “holes” in the gradient.

A cell has limited knowledge of the gradient, as it only perceives the chemotactic signal on the portion of Λ_2_ corresponding to the cell’s occupancy on Λ_1_. We define the vector 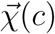 as the vector that spans from the cell’s center of mass to the center of mass of the perceived gradient. Copies of lattice sites are favoured when they align with the direction of the vector 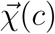, i.e. when there is a small angle *θ*_*c*_ between 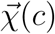 and the vector that spans from the center of mass of the cell to the lattice site into which copying is attempted (Fig. 1):

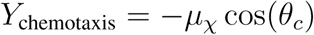

Where *µ*_*χ*_ is the maximal propensity to move along the perceived gradient. A uniform random *θ*_*c*_ ∈ [0, 2*π*[is chosen whenever 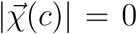, i.e. when, locally, there is no gradient (which may happen for very shallow gradients).

### 4.2 Evolutionary dynamics

A population of *N* cells undergoes the cell dynamics described above for the duration of a season, i.e. *τ*_*s*_ MCS. At the end of the season the evolutionary dynamics take place. The evolutionary dynamics are decoupled from the cell dynamics for the sake of simplicity, and consist of fitness evaluation, cell replication with mutation, and cell death to enforce constant population size.

#### Fitness evaluation

Fitness - i.e. the probability of replication - is calculated at the end of each season for each cell. We do not include any explicit advantage or disadvantage due to multicellularity, and instead assume that fitness is based only on individual properties of the cells. Therefore, any multicellular behaviour is entirely emergent in this simulation.

The fitness *F* (*c*) of a cell *c* ∈ *C* depends on the distance *d* = *d*(*c*) of the center of mass of a cell *c* from the peak of the gradient as a sigmoid function which is maximal when *d* = 0, and decreases rapidly for larger values of *d*:

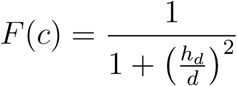

with *h*_*d*_ being the distance at which *F* (*c*) = 1/2.

#### Replication

For each cell *i* ∈ *C* with fitness *F*(*i*), the probability of replicating is P(cell *i* replicates) = *F*(*i*)/ ∑_*c* ∈ *C*_ *F* (*c*). We allow for *N* replication events, each calculated with the same probabilities, choosing only cells that were already present in the previous season (so not their offspring). Cells with larger fitness may be chosen multiple times for replication.

Each replicating cell divides along its short axis to create a daughter cell (see e.g. [37]), ensuring that related cells start close to each other at the beginning of the new season. One of the two cells (chosen randomly) is considered the mother cell, and can re-enter the competition for replication, the other cell may undergo mutations in their receptor and ligand composition. The bitstrings of the receptor and ligand may be modified with a per-position probability *µ*_R,L_. Mutations flip individual bits (from 0 to 1, and vice versa).

Because repeatedly halving a cell’s area would quickly lead to very small cells, we run a small number *η* of steps of the cell dynamics (without cell migration and chemotaxis) between two replication events that affect the same cell, so that cells can grow back to target size.

#### Death

After replication, there are 2*N* cells on the lattice. In order to restore the initial population size *N*, half of the cells are removed from the lattice at random. When the initial population size is restored, the season ends. The new season begins by randomly placing the peak of a new gradient at the mid-way point of a randomly chosen boundary (different from the previous one). The remaining cells will then undergo the cell dynamics for the following *τ*_*s*_ MCS.

## 5 Acknowledgment

We thank Paulien Hogeweg for constructive discussion. This research is supported by the Origins Center (NWA startimpuls). All authors declare no conflict of interest.

## 6 Author Contributions

ESC and RMAV conceived the study. ESC, RMAV, RMHM developed the model. ESC ran the simulations, analysed the data and wrote the first draft of the manuscript. ESC, RMAV, RMHM interpreted the results and wrote the final manuscript.

---

**Table 1:** Parameters.

| Parameter | explanation | Values |
| --- | --- | --- |
| $V^2$ | lattice size | $500 \times 500$ pix |
| $T$ | Boltzmann temperature | 16 AUE |
| $\lambda$ | cell stiffness | 5.0 AUE/pix<sup>2</sup> |
| $A_T$ | cell targetarea | 50 pix |
| *Cell adhesion* |  |  |
| $J_\alpha$ | minimum J value between cells | 4 AUE/[pix length] |
| $J'_\alpha$ | minimum J value between cell and medium | 8 AUE/[pix length] |
| $\nu$ | length of receptor and ligand bitstring | 24 bits |
| $\nu'$ | length ligand bitstring for medium adhesion | 6 bits |
| *Cell migration and chemotaxis* |  |  |
| $\mu_p$ | strength of persistent migration | 3.0 AUE |
| $\tau_p$ | duration of persistence vector | 50 MCS |
| $\mu_\chi$ | strength of chemotaxis | 1.0 AUE |
| $k_\chi$ | scaling factor chemokine gradient | 1.0 molecules/[pix length] |
| $p_{\chi=0}$ | probability of zero value ('hole') in gradient | $0.1 \text{ pix}^{-1}$ |
| *Evolution* |  |  |
| $N$ | population size | 200 cells |
| $\tau_s$ | duration of season | $5 \times 10^3 - 150 \times 10^3$ MCS |
| $h_d$ | distance from gradient peak where fitness is $\frac{1}{2}$ | 50 [pix length] |
| $\mu_{R,L}$ | receptor and ligand mutation probability | 0.01 per bit, per replication |

AUE: Arbitrary Units of Energy (see Hamiltonian in Model Section); pix: unit of area (one pixel of the lattice); pix length: unit of distance; MCS: Monte Carlo Step (unit of time).
<sup>468</sup> **Supplementary Material of:** <sup>469</sup> **Collective integration of spatial information drives the** <sup>470</sup> **evolution of multicellularity**

## Supplementary Material of: Collective integration of spatial information drives the evolution of multicellularity

### S1 Indistinguishable relative movement of cells with and without a chemotactic gradient

Here we investigate whether cells in a cluster move differently when they are performing chemotaxis or not. Fig. S1.1 shows the flow field around moving cells in a cluster with or without a gradient, as devised by [29]. In short, the flow field is calculated by taking each cell as a reference, and then rotating all other cells and their displacement vectors such that the reference cell displacement points to the right 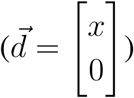. Then the rotated displacement vectors are summed in bins at defined points in the neighbourhood (using all the cells as a reference, and using different time points) to obtain the average displacements in the neighbourhood [29]. In this case, the flow field shows that the relative movement of cells in a cluster is the same whether there is a gradient or not.

**Figure S1.1:**
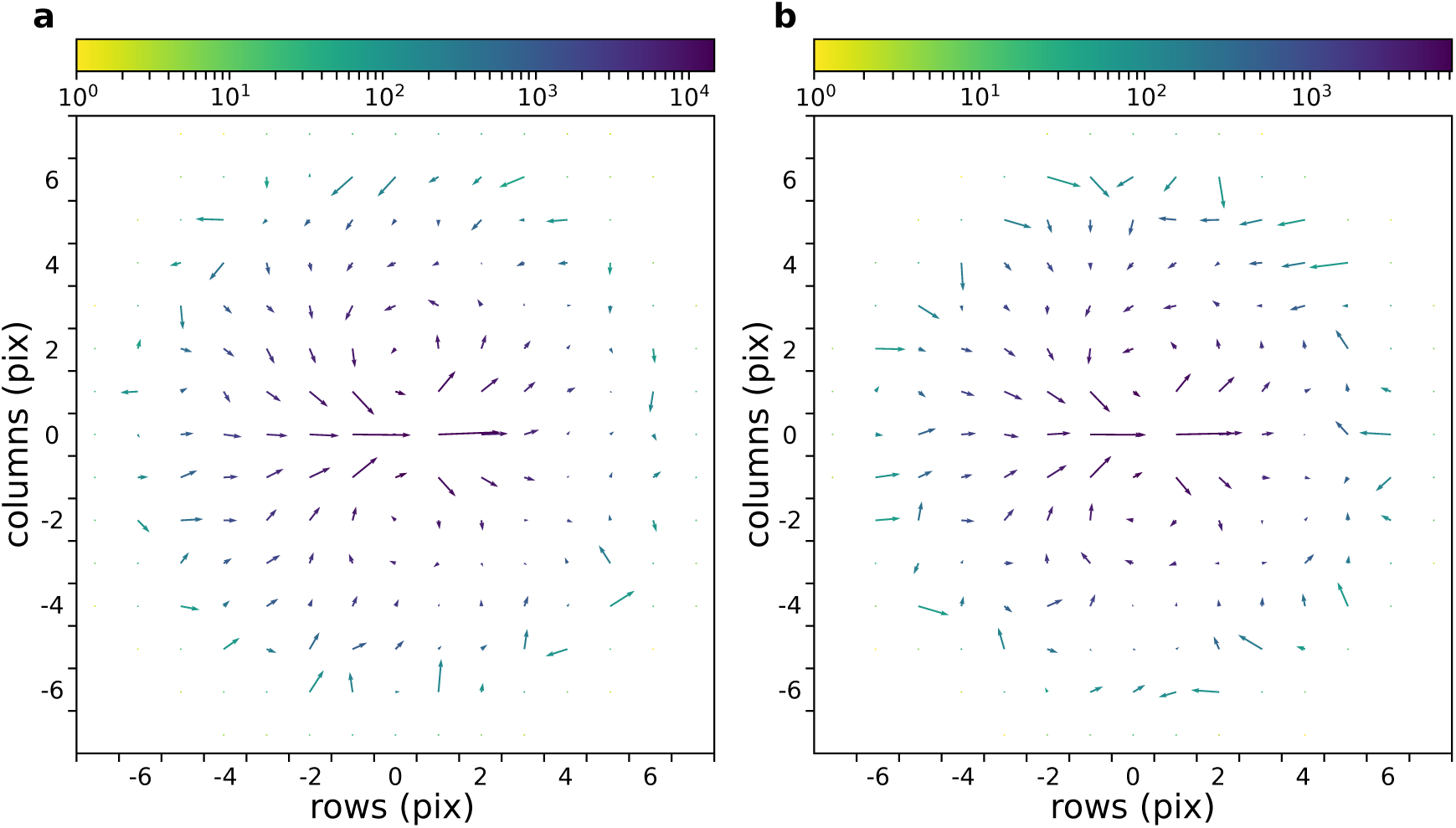
The flow field of a cluster of cells with and without gradient. **a** With chemokine gradient. **b** Without chemokine gradient. In both cases *N* = 50 cells with *γ* = 6 are placed at the center of the field (All other parameters as in main text).

### S2 Chemotaxis with short persistence of migration and small persistence strength

Fig. S2.2 shows that chemotaxis occurs in a rigid cluster of strongly adhering cells. The lower persistence strength reduces the number of changes in the relative position of cells within the cluster.

**Figure S2.2:**
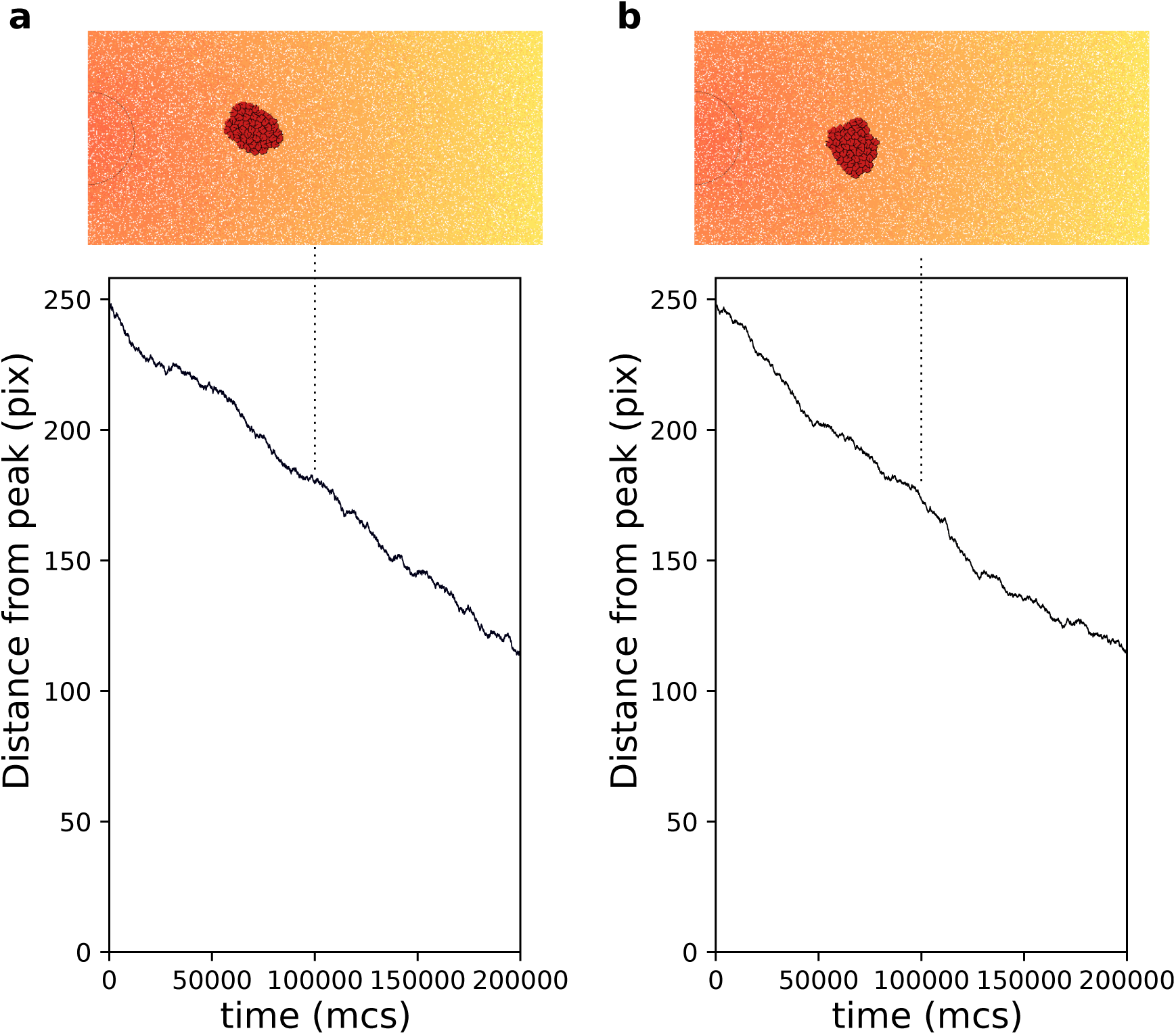
Chemotaxis of a rigid cluster. **a** *τ*_*p*_ = 5. **b** *µ*_*p*_ = 0.5. In both cases *N* = 50 cells with *γ* = 6 are placed on the right of the field and move towards higher concentration of the gradient (the semicircle indicates the resource location, where the gradient is highest. All other parameters as in main text).

### S3 Extraction of straight segments from cell tracks

For the contour plots in Fig.3 of the main text, we extracted straight segments of the cells’ trajectories, then measured the length of this segment and its angle with the direction of the source of the gradient. To identify these straight segments, we take increasingly longer intervals between the recorded cell positions, and measure how far the intermediate data points are positioned from the line spanning these two data points (Fig. S3.3A). As soon as one of the data points has a distance greater than a threshold, we stop extending the interval and continue from the cell position at which the chosen segment ends (the threshold value is set to 3 pixel lengths; this value is chosen because it is the largest integer smaller than the average cell radius, given a cell area = 50 pix). In figure S3.3B, the resulting segments are superimposed on cell position data from two simulations: one with a single cell and one with a cluster of adhering cells. While the overlap between the segment and the track itself varies, the length and orientation of the straight parts of the track are generally well-preserved in the segments.

**Figure S3.3:**
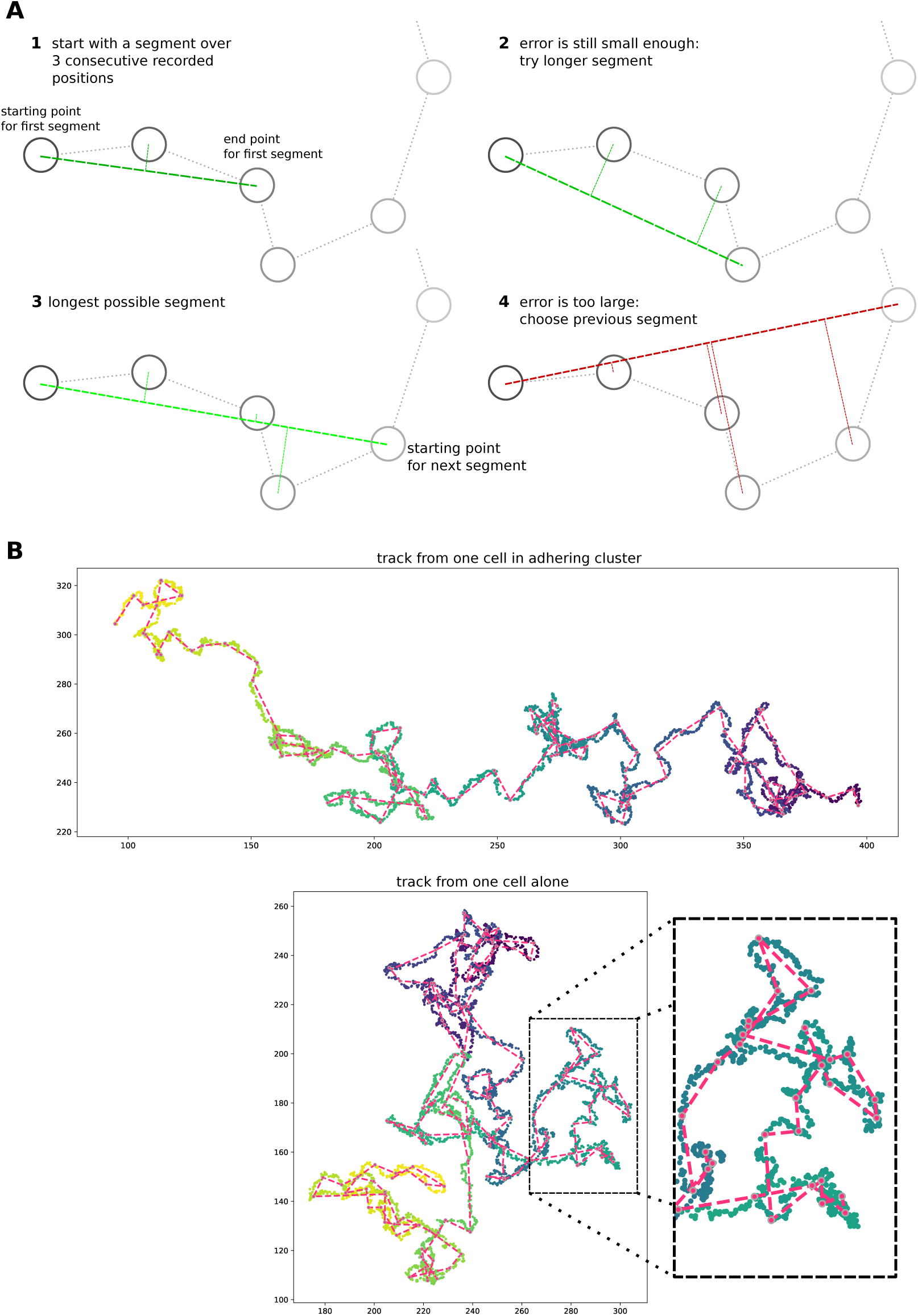
Simple algorithm for segment extraction. **a)** Visual explanation of the algorithm, with a cartoon representation of a cell track with cell positions recorded at regular time intervals. Images 1-4 represent subsequent stages of the algorithm. For 1-3, the maximum distance of intermediate cell positions is still small enough, while for the segment in image 4 two intermediate positions are too far away. So the segment in image 3 will be used in the analysis, and we will start the algorithm from the fourth data point. **b)** Two cell tracks from simulations, with the extracted segments superimposed in red.

### S4 Chemotaxis of cells with different *A*_*T*_

We explored the behaviour of different cell sizes and cell number by running simulations where the total area of the cells is kept constant, *NA*_*T*_ = 5000. We expect that large cells move with greater persistence towards the peak of the gradient than small cells, because they perceive a larger portion of the gradient, thus averaging out noise. Indeed, Fig. S4.4 shows that larger cells perform chemotaxis more efficiently than smaller cells, given the same chemotactic gradient.

**Figure S4.4:**
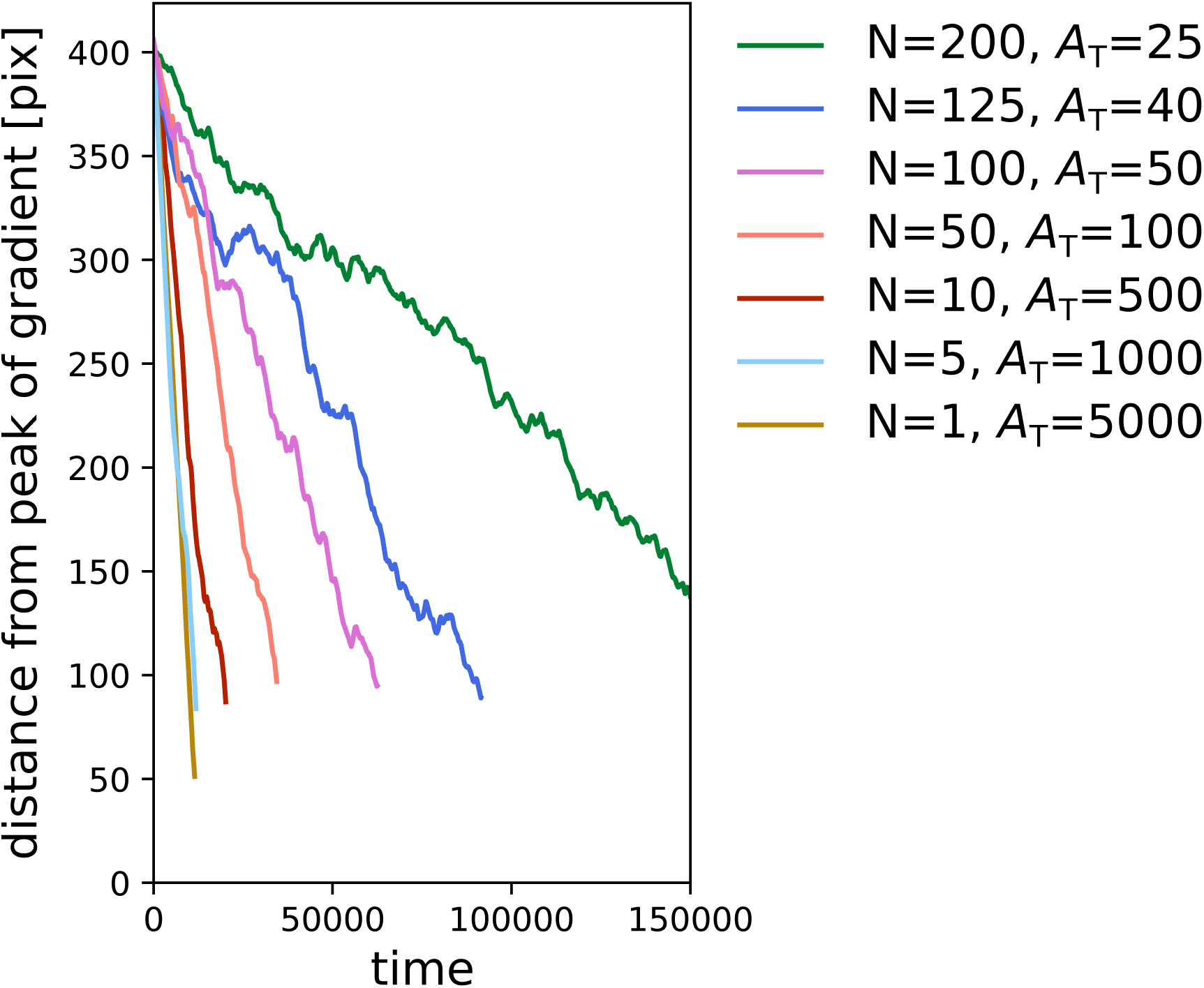
Large cells perform chemotaxis more efficiently than clusters of small cells. Each line corresponds to one simulation with a given combination of number of cells *N* and cell size *A*_*T*_, and shows the distance of the centre of mass of the cluster of cells from the peak of the gradient over time. We kept the total volume of the cells constant in all simulations (i.e. *NA*_*T*_ = 5000). All other parameters (including the chemotactic signal) are the same as in main text.

### S5 Supplementary videos

1. Migrating cluster of adhering cells. Cell colour indicates the direction of migration, to emphasize the dynamics within the cluster.
2. The same cluster of adhering cells. All cells have the same colour to show how the migration of the cluster as a whole resembles that of an amoeba.
3. Video of an evolutionary simulation, starting with neutrally adhering cells (*γ* = 0). The season changes every 100 * 10^3^ MCS.

